# Further varieties of ancient endogenous retrovirus in human DNA

**DOI:** 10.1101/2024.12.11.627920

**Authors:** Martin C. Frith

**Affiliations:** Department of Computational Biology and Medical Sciences, University of Tokyo; Artificial Intelligence Research Center, AIST; AIST-Waseda University Computational Bio Big-Data Open Innovation Laboratory

## Abstract

A retrovirus inserts its genome into the DNA of a cell, occasionally a germ-line cell that gives rise to descendants of the host organism: it is then called an endogenous retrovirus (ERV). The human genome contains relics from many kinds of ancient ERV. Some relics contributed new genes and regulatory elements.

This study finds further kinds of ancient ERV, in the thoroughly-studied human genome version hg38: ERV-Hako, ERV-Saru, ERV-Hou, ERV-Han, and ERV-Goku. It also finds many relics of ERV-V, previously known from just two copies on chromosome 19 with placental genes. It finds a type of ERV flanked by MER41E long terminal repeats (LTRs), with surprisingly little similarity to the known MER41 ERV. ERV-Hako has subtypes that contain sequence from host genes *SUSD6* and *SPHKAP* : the *SUSD6* variant was transferred between catarrhine and platyrrhine primates. A retrovirus uses tRNA to prime reverse transcription: Hako is the only human ERV relic that used tRNA-Trp (tryptophan, symbol W), and HERV-W is misnamed because it used tRNA-Arg, based on the Genomic tRNA Database. One ERV-Saru LTR is the previously-described enhancer of *AIM2* in innate immunity. This study contributes to understanding primate ERV history, but also shows that related ERVs can have drastic differences, challenging the goal of clearly annotating all ERV relics in genomes.

## Introduction

After a retrovirus enters a cell, it reverse-transcribes its RNA genome into DNA and inserts it into the cell’s genome, where it is called a provirus. Occasionally this happens in a germ-line cell, and the provirus is inherited by the host animal’s descendants: it is then called an endogenous retrovirus (ERV). Like any mutation, the ERV may spread by parent-to-child inheritance until it becomes fixed in the host species’ genome, or it may be lost from the population.

Vertebrate genomes have many relics of ancient ERVs: they comprise about 8% of the human genome. They are “fossils” that tell us about ancient virus evolution. Some relics evolved into vertebrate genes and regulatory elements. Typically, one kind of ERV has many relics scattered through the genome. This is due to germ-line re-infection, alternatively, some ERVs evolved to replicate within a cell as retrotransposons [1]. Annotation of ERV relics in genomes (along with other types of repeated element) is often done with the RepeatMasker software [2] and the Dfam database of repeat elements [3].

A retroviral provirus has a duplicated segment at its two ends, termed long terminal repeats (LTRs). Between them is an interior region that encodes gag, pol, and env proteins. Most ERV relics, however, are solo LTRs. These presumably arise from homologous recombination between the two LTRs of an ERV, which removes the interior and leaves a single LTR.

A virus may undergo mutation that destroys viral functioning: such a “defective” virus can nevertheless propagate non-autonomously, using molecules from a “helper” virus. A retroviral genome can get joined to a host gene: this can produce new kinds of retrovirus (usually defective) that contain parts of host genes [4]. Such host gene capture was found in crocodilian ERV relics [5]. Koala retrovirus (KoRV) is currently endogenizing in koalas: it has defective forms known as RecKoRV. Interestingly, some koala populations have RecKoRV only, though presumably KoRV must have been there in the past as a helper virus [6].

The retrovirus family *Retroviridae* (in class *Revtraviricetes*) is classified into genera including *Alpharetrovirus, Betaretrovirus, Gammaretrovirus, Deltaretrovirus, Epsilonretrovirus*, and others. Retrovirus classification should be integrated with ERV relics. ERVs have been classified, however, into groups I (related to *Gamma*- and *Epsilonretrovirus*), II (related to *Alpha*- and *Betaretrovirus*), III, and more.

Finer classification of ERVs is complex. Vargiu et al. classified human ERV relics into 39 clades plus 31 less well-defined groups with many mosaic forms [7]. They focused on relatively complete relics of ERVs that were not drastically defective.

Dfam version 3.8 has 566 types of human ERV relic (41% of all human repeat types), split into 436 LTRs and 130 interior regions. Many of these ERV types seem to lack detailed publications explaining them, and have cryptic names. This hinders learning, and understanding in adjacent fields like genomics. Names like MER41E are hard to remember, and could be any kind of repeat (MEdium Reiteration). Non-specialists often confuse “a mobile element that has LTRs” with “an LTR”. Some RepeatMasker names (e.g. HERV17) differ from the usual name among ERV researchers (HERV-W). A reasonable suggestion is to name ERV types as ERV-*name* (e.g. ERV-W), where the “ERV-” prefix can be dropped when clear from context [8].

In any case, the types of ancient ERV (and other mobile element) in human DNA have been thoroughly collected. Very ancient and degraded (e.g. Paleozoic) relics are arbitrarily hard to find [9], but maybe all younger types have been found? To investigate this, the present study sought unknown types of mobile element relic, in the thoroughly-studied human genome version hg38. It found new types of not-so-ancient ERV (Table 1). It found other elements not in Dfam that turn out to be previously known: Alu-related ASR and CAS [10], an SVA retrotransposon precursor [11], and an ERV-T1 LTR [12]. All these results have been submitted to Dfam.

**Table 1.** ERV sequences reconstructed in this study.

| Type | Part | Length | ERV group | TSD length (bp) | tRNA primer | Hosts | Approx. human copies<sup>a</sup> |
| --- | --- | --- | --- | --- | --- | --- | --- |
| ERV-Hako | LTR | 401 | II | 5-6 | Trp | Simiiformes | 47 |
|  | interior | 7085 |  |  |  |  | 16 |
| ERV-V | LTR | 721 | I | 4 | Val, His | Simiiformes | 89 |
| ERV-Saru | LTR | 636 | I | 4 | ? | Simiiformes | 14 |
| ERV-Hou | LTR | 681 | I | 4 | His | Simiiformes | 36 |
| ERV-Han | LTR | 698 | I | 4 | ? | Simiiformes | 60 |
| ERV-Goku | LTR | 545 | I | 4 | Arg | Primates | 86 |
| ERV-MER41E | interior | 4389 | I | 4 | Arg | Simiiformes | 15 |

<sup>a</sup> Per haploid human genome (hg38). A copy that is fragmented (e.g. by transposon insertions) is counted as 1. An incomplete copy is counted as 1.

## Methods

This study sprang from finding human genome regions homologous to transposable element proteins [9]. Some regions homologous to ERV proteins lack clear RepeatMasker annotation, indicating an unknown type of ERV (Fig. 1).

**Figure 1:**
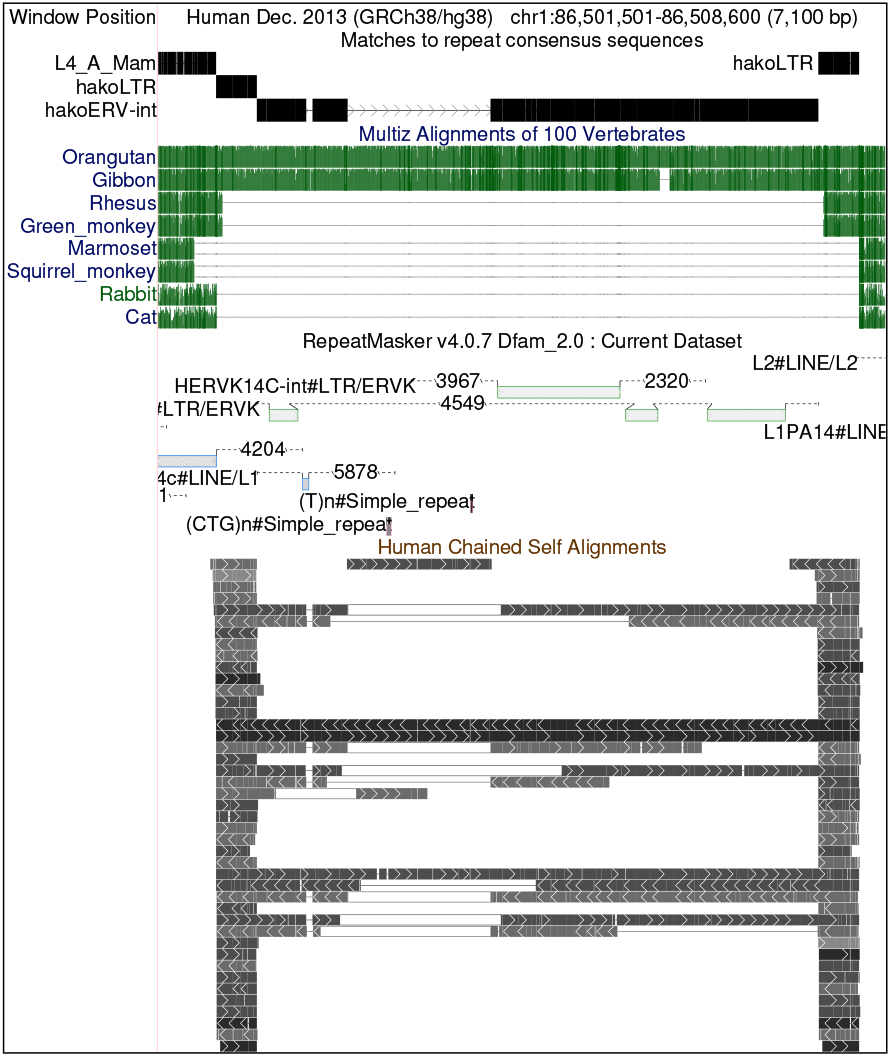
An ERV-Hako relic in human chromosome 1. hakoERV-int refers to the the interior region, which is flanked by LTRs. Screenshot from http://genome.ucsc.edu [16].

Next, new types of mobile element were sought from gaps in genome alignments. For example, if DNA sequence is present in apes but absent in other primates, it indicates an insertion in a common ancestor of apes. This may be a new type of element if a match to known elements is absent, partial, or of unexpected age, for example, an old type of element for a young insertion.

Also, biased coverage of Dfam elements was examined. For example, if an LTR’s left half occurs several times in the genome, it may indicate a new type of LTR whose left half only is similar to the known type.

The edges of a relic were inferred from genome alignment gaps, and target site duplications (TSDs) (Fig. S1). Often, mobile element insertion duplicates some DNA at the insertion site (4–6 bp for a retrovirus) to both sides of the insert. Sometimes, an element inserts inside an older element: then its edges can be inferred from the gap in the older element. Finally, ERVs start with tg and end with ca.

For each new type of element, several instances were fed to Refiner, which aligns them and makes a consensus sequence [13]. These consensus sequences were searched in mammal genomes (Table S1), by adding them to Dfam’s consensus sequences for that genome, then using “split-alignment” [14]. This optimizes alignments where each genome base-pair matches at most one base-pair from all consensus sequences, thus adjudicating between similar consensus sequences [15]. Method details are in the Supplement.

## Results

### ERV-Hako

ERV-Hako is a group II ERV, with a tendency to contain parts of host genes. (Hako is Japanese for box/container). An example is shown in Fig. 1. The genome alignments (green) show this ERV was inserted in a common ancestor of catarrhines (apes and Old World monkeys), after their last common ancestor with platyrrhines (New World monkeys, e.g. marmoset and squirrel monkey). It was reduced to a solo LTR in Old World monkeys (rhesus and green monkey), but not in apes (gibbon, orangutan, human).

The RepeatMasker annotation (Fig. 1) shows patchy similarity to known ERV-K sequences, but doesn’t recognize the LTRs. The human self alignments show that the LTRs are repeated many times in the genome, and the interior is occasionally repeated. The Hako consensus sequence has regions homologous to known ERV proteins (Fig. S2), and Hako insertions produced target site duplications of 5 or 6 bp (Fig. S1).

This ERV relic contains a segment that doesn’t match the Hako consensus sequence (Fig. 1 arrowheads). This segment matches parts of the 3^*′*^-UTR (untranslated region) of a gene: *SUSD6* (sushi domain containing 6) (Fig. 2). Other Hakos on chromosomes 3 and X also have this *SUSD6* insert. Another Hako on chromosome 1 has part of the 3^*′*^-UTR of *SPHKAP* (SPHK1 interactor, AKAP domain containing) (Fig. S3). This is similar to a retrovirus carrying a cellular gene [4].

**Figure 2:**
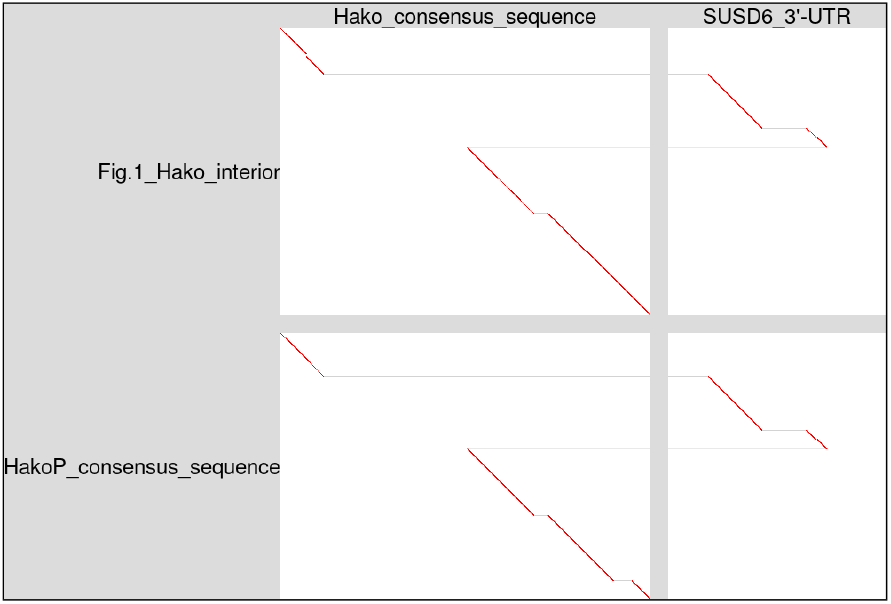
The Hako relic from Fig. 1 (vertical, top) mostly matches the Hako conensus sequence (horizontal, left), but partly matches the 3^*′*^-UTR of *SUSD6* (horizontal, right). The HakoP consensus sequence (vertical, bottom) is similar. The diagonal lines show which parts of the vertical sequences are similar to which parts of the horizontal sequences.

There is a similar type of ERV, AloPal-5.1366, in the New World howler monkey *Alouatta palliata* [3]. Here it is named ERV-HakoP (for platyrrhine). Remarkably, the HakoP consensus sequence has the same *SUSD6* insertion (Fig. 2). The ERV in Fig. 1 is intermediate between Hako and HakoP: its LTRs are much more similar to Hako (Fig. S4). No HakoP LTRs were found in non-platyrrhines, but platyrrhines have both HakoP and Hako-like LTRs. Alignments between platyrrhine genomes (Table S2) show that most of these insertions are common to diverse platyrrhines (howler monkey, marmoset, squirrel monkey, titi). Thus, they were inserted before the last common ancestor of platyrrhines *∼* 21 million years ago (mya) [17]. Full-length Hako ERVs were not found in platyrrhines, but an intermediate form was found (Fig. S5).

Most Hako insertions are common to catarrhines, or common to platyrrhines, but not common to both. However, two Hako relics are shared by catarrhines and platyrrhines (Fig. S6). This suggests ERV-Hako entered the genomes shortly before the end of gene flow between diverging catarrhines and platyrrhines.

Retrovirus reverse transcription is primed from the 3^*′*^-end of tRNA, base-paired with a PBS (primer binding site) at the 5^*′*^-edge of the interior region. The type of tRNA was inferred by comparing ERV sequences to human mature tRNAs in GtRNAdb release 21 [18]. This indicated that Hako and HakoP used tRNA-Trp (Fig. 3), whereas most group II human ERV relics used tRNA-Lys. Remarkably, Hako is the only type of human ERV relic that used tRNA-Trp, according to GtRNAdb. Another ERV was named HERV-W because it was thought to use tRNA-Trp. Vargiu et al. mentioned ambiguity between the tRNA-Trp sequence “tggcgac-cacgaagggac” and tRNA-Arg in the Leipzig tRNA database [7]. According to GtRNAdb, however, this sequence is tRNA-Arg. (At the time of writing this, the Leipzig database is not available.)

**Figure 3:**
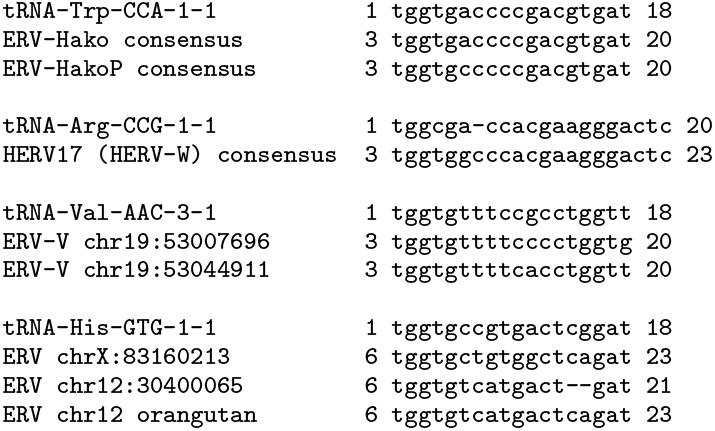
Alignments between human mature tRNAs from GtRNAdb (reverse-complemented), and ERV interior sequences.

### ERV-V

ERV-V is a group I ERV, which infected a common ancestor of simians, and has two copies in the human genome, which are near neighbors in chromosome 19 [19, 20]. A third copy lies between them in non-hominines. Remarkably, they encode proteins derived from env, gag, and an unusual pre-gag gene, which are expressed in placenta and exhibit evolutionary patterns suggesting they contribute to the animal’s fitness [19–21]. Previous studies found no other copies or solo LTRs [19, 21], though Boso et al. found similar gag and pre-gag sequences in five other human ERV relics [20].

The present study reconstructed a consensus sequence that matches the ERV-V LTRs and occurs about 90 times in the human genome, mostly as solo LTRs. Parts of this LTR are similar to other LTRs in Dfam (Fig. S7). One of the five ERVs found by Boso et al. in chromosome X has this LTR at it’s 5^*′*^-end. (The 3^*′*^-end is missing.) Another ERV relic in chromosome 12 has this LTR at both ends (Fig. S8). These two ERVs have partial similarity to the chromosome 19 ERV-Vs (Fig. 4), and to each other (Fig. S9). They have fragmentary matches to gag, pol, and env genes (Fig. S10). Surprisingly, they were primed by tRNA-His, whereas the chromosome 19 ERV-Vs were primed by tRNA-Val (Fig. 3).

**Figure 4:**
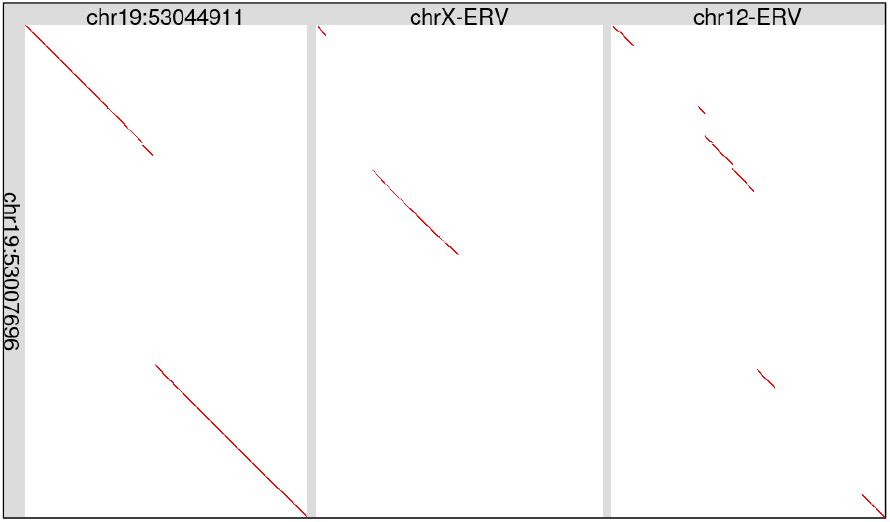
Two ERV relics, in human chromosomes 12 and X, are partially similar to ERV-V in chromosome 19. The two chromosome 19 ERVs are similar to each other apart from one gap. This figure just shows the interior ERV regions, excluding the LTRs. The ERVs are shown after removing retrotransposons (e.g. Alu) inserted within them.

### ERV-Saru

ERV-Saru entered the genome in a common ancestor of simians ≳43 mya, after their last common ancestor with other primates *∼* 69 mya. (Saru is Japanese for monkey.) It survives only as solo LTRs. It is similar to other types of LTR: most of it is similar to MER41G (59% identity), but its last 100 bp are more similar to LTR43B and its first 100 bp are not similar to any known LTR (Fig. S11). Surprisingly, MER41G is classified as *Gammaretrovirus* and LTR43B as *Epsilonretrovirus* [22]. Remarkably, one Saru LTR regulates innate immunity: it is an interferon-induced enhancer of *AIM2* (Absent in Melanoma 2) [23]. It was thought to be a MER41 element, but is unambiguously a full-length Saru LTR.

### ERV-Hou

ERV-Hou also entered the genome in a common ancestor of simians, after their last common ancestor with other primates. (Hou is Chinese for monkey.) It survives mostly as solo LTRs, but there are two interior ERV regions, in chromsomes 1 and 22. These interiors are homologous only in their 3^*′*^ parts (Fig. 5). Some parts of these interiors are homologous to other types of ERV (Fig. S12), which are classified as *Gammaretrovirus* [22].

**Figure 5:**
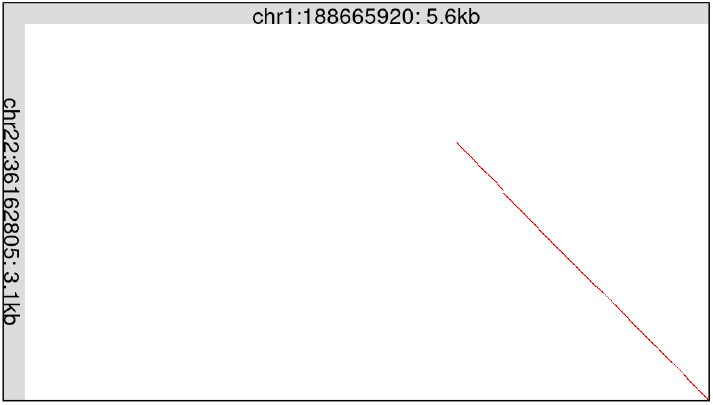
Two human ERV relics with Hou LTRs are only partially similar. This figure just shows the interior regions, between the LTRs. The chromosome 1 ERV (horizontal) is shown after deleting a recentlyinserted LINE fragment.

Only a small fragment of the Hou LTR is similar to other known LTR types (Fig. S13). Other fragments of the Hou LTR, however, have repeated homologs in diverse mammals (Fig. S13). This suggests that those mammals harbor unknown types of LTR partially homologous to Hou LTR. Near-full-length homologs of Hou LTR appear in Cetruminantia (ruminants, hippos and whales) (Fig. S14).

### ERV-Han

ERV-Han survives only as solo LTRs (Fig. 6). It entered the genome in a common ancestor of simians, after their last common ancestor with other primates. This LTR’s 3^*′*^ half is similar to LTR43B (*Epsilonretrovirus*), it has a middle region similar to the ERV-Goku LTR, and its 5^*′*^ region is not similar to any other repeat element (Fig. S15).

**Figure 6:**
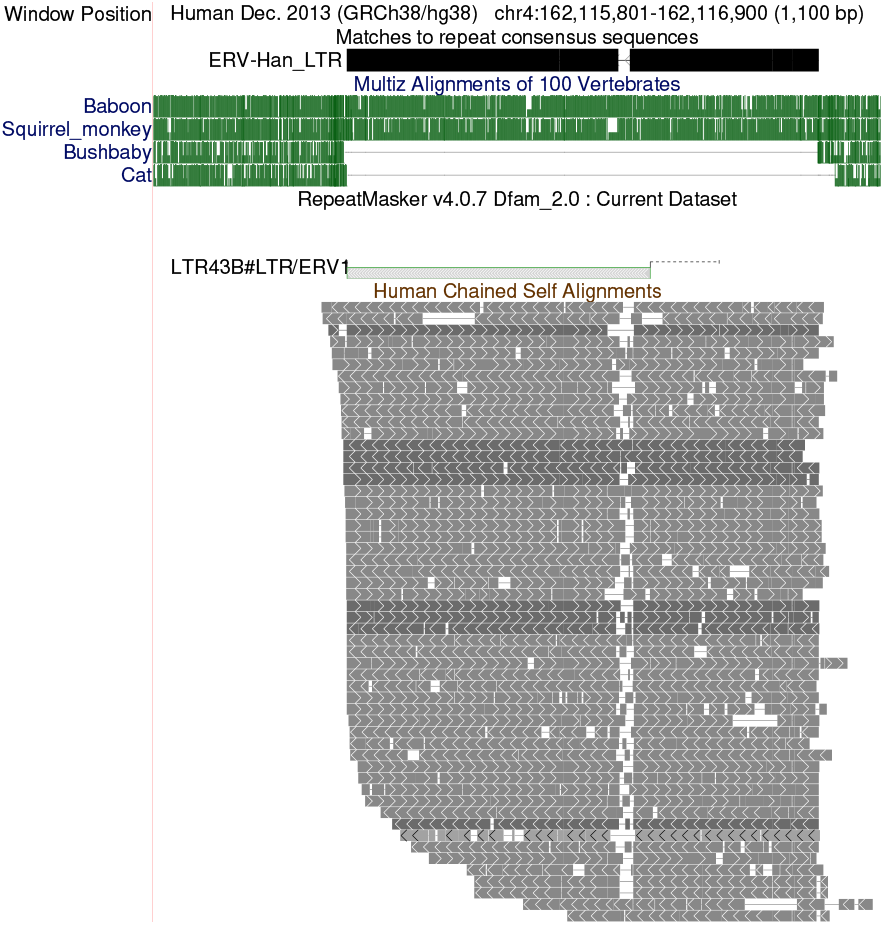
An ERV-Han solo LTR in human chromosome 4. The self alignments indicate that there are similar sequences elsewhere in the genome.

### ERV-Goku

ERV-Goku also entered the genome before the last common ancestor of simians, after their last common ancestor with other primates. (Goku is a simian story character.) It survives mostly as solo LTRs, but there are four interior regions in hg38, on chromosomes 3, 13, 19, X, and a fifth fragment on chromosome 10. Part of the LTR is similar to LTR35B (*Gammaretrovirus*), and part is similar to the ERV-Han LTR (Fig. S16). The interior regions have patchy similarity to other ERVs classified as *Gammaretrovirus* (Fig. S17). They also have full-length homology to gag, pol, and env proteins (Fig. S17), suggesting that ERV-Goku may have been autonomous. Surprisingly, ERV-Goku also appears in non-simian primate genomes (Fig. S18). Goku LTR relics are shared by mouse lemur, slow loris, and bushbaby: they entered the genome before the last common ancestor of strepsirrhines *∼* 59 mya. Goku LTR relics are also in tarsiers. Thus, all primates have Goku relics, which may have entered their genomes around the same time. Goku was not found in colugo (the closest relative to primates) or other mammals (Table S1).

### ERV-MER41E

A MER41 LTR was described in [24]. Today, Dfam has LTR subtypes MER41A, B, C, D, E, and G, and one interior sequence MER41-int. They are all classified as *Gammaretrovirus*, though MER41C is surprisingly in a separate ERV sub-group [22].

Some MER41 LTR relics bind to STAT1 transcription factors and regulate innate immunity [23, 25]. Others overlap placental enhancers and bind to SRF and other transcription factors [26, 27]. One MER41 evolved to produce the noncoding RNA *BANCR*, which plays a role in heart development [28].

This study reconstructed an interior sequence that appears between MER41E LTRs. Remarkably, just its final 108 bp are similar to MER41-int. Instead, it has patchy similarity to other ERV sequences in Dfam (Fig. S19). It matches gag, pol, and env proteins, but has a large deletion overlapping pol and env, implying it was non-autonomous (Fig. S19).

Genome alignments suggest that all MER41 subtypes entered the genome in a common ancestor of simians, after its last common ancestor with other primates. However, MER41-like ERVs colonized other mammals independently [23]. These MER41-like ERVs are similar to parts of the MER41 ERVs (Fig. S20). Interestingly, tarsier has LTR relics with full-length similarity to MER41E and MER41G (Fig. S21).

## Discussion

This study found new types of not-so-ancient ERV in the thoroughly-studied human genome version hg38: thus not all types were found already. This contributes to understanding our genome’s history. For example, the *AIM2* enhancer comes from a specific low-copy ERV type ERV-Saru, and some half LTR43B elements are actually half of ERV-Han LTRs.

It was hoped this study might get close to finding all types of not-too-ancient mobile element. In parallel, however, Takeda et al. found two new types of LTR (present in hg38), with no overlap to those found here [29]. This suggests we are still not finished.

An appealing goal is to reconstruct all types of ancient mobile element, so that we can match the genome against them and clearly annotate all relics. This may be unachievable due to drastic variation among related elements, especially ERVs. For example, there are four ERV-V interiors in hg38: two are tandem duplicates, and the other two are so different that it seems hard or inappropriate to reconstruct one original sequence.

It is interesting that similar ERV insertions are in different primate lineages though they seem to be younger than the last common ancestor of those lineages. This could be explained by inter-species viral transmission, or interbreeding and gene flow. A low level of gene flow from lineage A to B might not get fixed in the B genome, leaving no trace, except that active mobile elements could be transmitted. By one of these mechanisms, an ERV-Hako containing part of a host gene was transmitted between catarrhines and platyrrhines, indicating that host DNA can be transferred between lineages. Also, ERVs in one lineage can be better understood by studying other lineages. For example, platyrrhines have only defective Hakos, like koalas with only RecKoRV: catarrhines reveal the ancestral ERV-Hako.

## Acknowledgments

Arian Smit co-authored the MER41E interior region reconstruction.

