## Supplement for "Further varieties of ancient endogenous retrovirus in human DNA"

Martin C. Frith

December 11, 2024

### Methods

#### Matching genomes to repeat consensus sequences

Each genome (Table S1) was matched to repeat consensus sequences as follows. The new consensus sequences (Table 1), plus HakoP and all MER41 sequences from Dfam (version 3.8), were added to all Dfam consensus sequences for that genome. The matching was done with LAST version 1592 (`https://gitlab.com/mcfrith/last`). First, an index (named repDB) was made for the repeat sequences:

```
lastdb -uMAM8 -S2 repDB repeats.fasta
```

- `-uMAM8` makes the matching more sensitive but slow [1].
- `-S2` indexes both DNA strands of the repeats (so we can compare to just one strand of the genome).

Then, rates of matches, mismatches and gaps between repeats and genome were found [2]:

```
last-train -P8 --revsym -X1 --pid=70 repDB genome.fasta > rep.train
```

- `-P8` makes it faster by using 8 threads, with no effect on results.
- `--revsym` forces strand-symmetric rates, e.g. `a`→`g` equals `t`→`c`.
- `-X1` treats matches to unknown `n` bases in the consensus sequences as neutral instead of disfavored.
- `--pid=70` makes the training ignore alignments with >70% identity. This makes it better at finding old elements, but worse at finding young elements. 70 seems to be a good compromise.

Finally, the genome was matched to the consensus sequences:

```
lastal -P8 -D1e7 -p rep.train --split repDB genome.fasta | last-postmask > out.maf
```

- `--split` selects the split-alignment algorithm, which makes each genome base-pair align to at most one base-pair from all consensus sequences [3].
- `-D1e7` gets alignments with similarity score expected by chance between random sequences at most once per $10^7$ genome base-pairs versus *all* the consensus sequences. So we expect about 300 spurious matches to the human genome, which is a tiny fraction of the total matches.
- `last-postmask` removes alignments caused by simple sequence such as `attattattattattatttttataat`.

For display in the genome browser, the alignments were converted to PSL format:

```
maf-convert -s2 -j10000 psl out.maf > out.psl
```

#### DNA-versus-protein sequence alignments

Some of the following figures show alignments between DNA and protein sequences. They were made like this:

```
lastdb -q -c protDB proteins.fasta
lastal -m1000 -p te.train protDB dna.fasta
```

`te.train` is a file with rates of matches, mismatches, gaps, and frame-shifts, from a previous study [4], available in the hg38 directory of https://github.com/mcfrith/protein-fossils.

#### Genome-to-genome alignments

Pairs of genomes (Table S2) were aligned like this:

```
lastdb -P8 -c -uRY4 myDB genome1.fasta

last-train -P8 --revsym -C2 myDB genome2.fasta > my.train

lastal -P8 -D1e9 -C2 --split -p my.train myDB genome2.fasta |
last-split -r | maf-linked - > out.maf
```

For alignments of human to non-simians, `RY4` was replaced by `MAM4`, which makes it more sensitive but slow.

#### Alignments between ERV DNA sequences

Alignments between ERV DNA sequences (e.g. Fig. 4, 5) were found with LAST version 1607:

```
lastdb -c horiDB horizontalSequences.fasta
lastal horiDB verticalSequences.fasta
```

For Fig. 2, `--split` was added to the `lastal` command.

**Table S1:** Genome versions used in this study

| Animal | Clade | Scientific name | Genome |
|---|---|---|---|
| human | Catarrhini | *Homo sapiens* | hg38_no_alt_analysis_set |
| chimpanzee | Catarrhini | *Pan troglodytes* | GCF_028858775.1 |
| orangutan | Catarrhini | *Pongo abelii* | GCF_028885655.1 |
| gibbon | Catarrhini | *Nomascus leucogenys* | GCF_006542625.1 |
| rhesus macaque | Catarrhini | *Macaca mulatta* | GCF_003339765.1 |
| snub-nosed monkey | Catarrhini | *Rhinopithecus roxellana* | GCF_007565055.1 |
| marmoset | Platyrrhini | *Callithrix jacchus* | GCF_011100555.1 |
| squirrel monkey | Platyrrhini | *Saimiri boliviensis* | GCF_016699345.2 |
| howler monkey | Platyrrhini | *Alouatta palliata* | GCA_004027835.1 |
| titi | Platyrrhini | *Cheracebus lugens* | GCA_963574535.1 |
| tarsier | Haplorhini | *Carlito syrichta* | GCF_000164805.1 |
| mouse lemur | Strepsirrhini | *Microcebus murinus* | GCF_000165445.2 |
| slow loris | Strepsirrhini | *Nycticebus coucang* | GCF_027406575.1 |
| bushbaby | Strepsirrhini | *Otolemur garnettii* | GCF_000181295.1 |
| colugo | Dermoptera | *Galeopterus variegatus* | GCF_000696425.1 |
| treeshrew | Scandentia | *Tupaia tana* | GCA_026018925.1 |
| rabbit | Lagomorpha | *Oryctolagus cuniculus* | GCF_009806435.1 |
| marmot | Rodentia | *Marmota marmota* | GCF_001458135.2 |
| mouse | Rodentia | *Mus musculus* | GCF_000001635.27 |
| dolphin | Artiodactyla | *Tursiops truncatus* | GCF_011762595.1 |
| minke whale | Artiodactyla | *Balaenoptera acutorostrata* | GCF_949987535.1 |
| hippo | Artiodactyla | *Hippopotamus amphibius* | GCF_030028045.1 |
| cow | Artiodactyla | *Bos taurus* | GCF_002263795.3 |
| mouse-deer | Artiodactyla | *Tragulus kanchil* | GCA_022376925.1 |
| pig | Artiodactyla | *Sus scrofa* | GCF_000003025.6 |
| alpaca | Artiodactyla | *Vicugna pacos* | GCF_000164845.4 |
| horse | Perissodactyla | *Equus caballus* | GCF_002863925.1 |
| dog | Carnivora | *Canis familiaris* | GCF_011100685.1 |
| fruit bat | Chiroptera | *Pteropus vampyrus* | GCF_000151845.1 |
| elephant | Afrotheria | *Loxodonta africana* | GCF_000001905.1 |
| armadillo | Xenarthra | *Dasypus novemcinctus* | GCF_030445035.1 |

**Table S2:** Pairs of genomes that were aligned to each other

| genome1 | genome2 |
|---|---|
| human | chimpanzee |
| human | orangutan |
| human | gibbon |
| human | rhesus macaque |
| human | snub-nosed monkey |
| human | marmoset |
| human | squirrel monkey |
| human | tarsier |
| human | mouse lemur |
| human | slow loris |
| human | bushbaby |
| marmoset | squirrel monkey |
| marmoset | howler monkey |
| marmoset | titi |
| bushbaby | slow loris |
| bushbaby | mouse lemur |

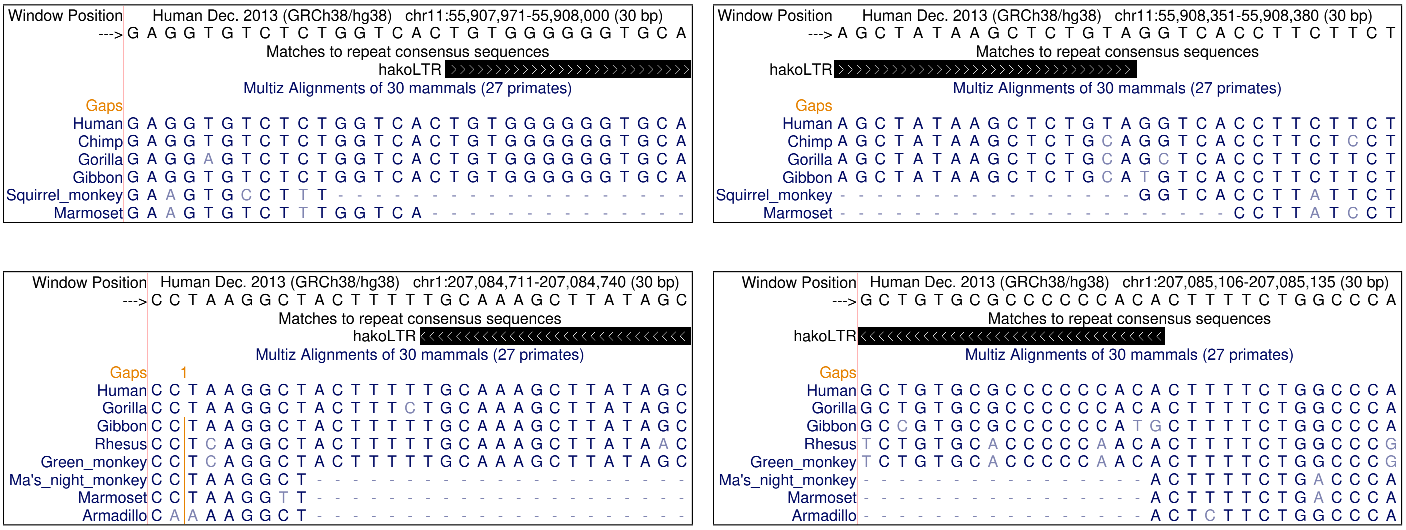


**Figure S1:** Above: The left and right ends of one ERV-Hako solo LTR in human chromosome 11, showing target site duplication `ggtcac`. Below: The left and right ends of one ERV-Hako solo LTR in human chromosome 1, showing target site duplication `ctttt`.

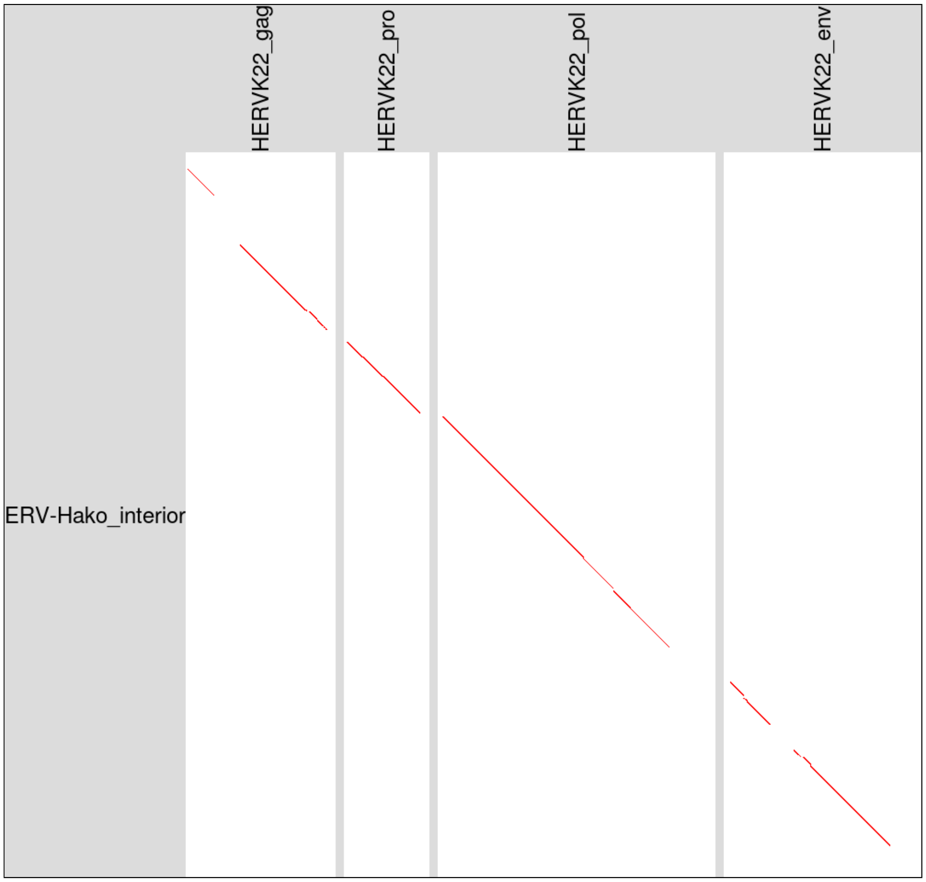


**Figure S2:** Homology between the ERV-Hako DNA consensus sequence (vertical) and retroviral proteins (horizontal). This figure was made by DNA-versus-protein alignment with LAST. The proteins are from RepeatMasker's file of protein sequences.

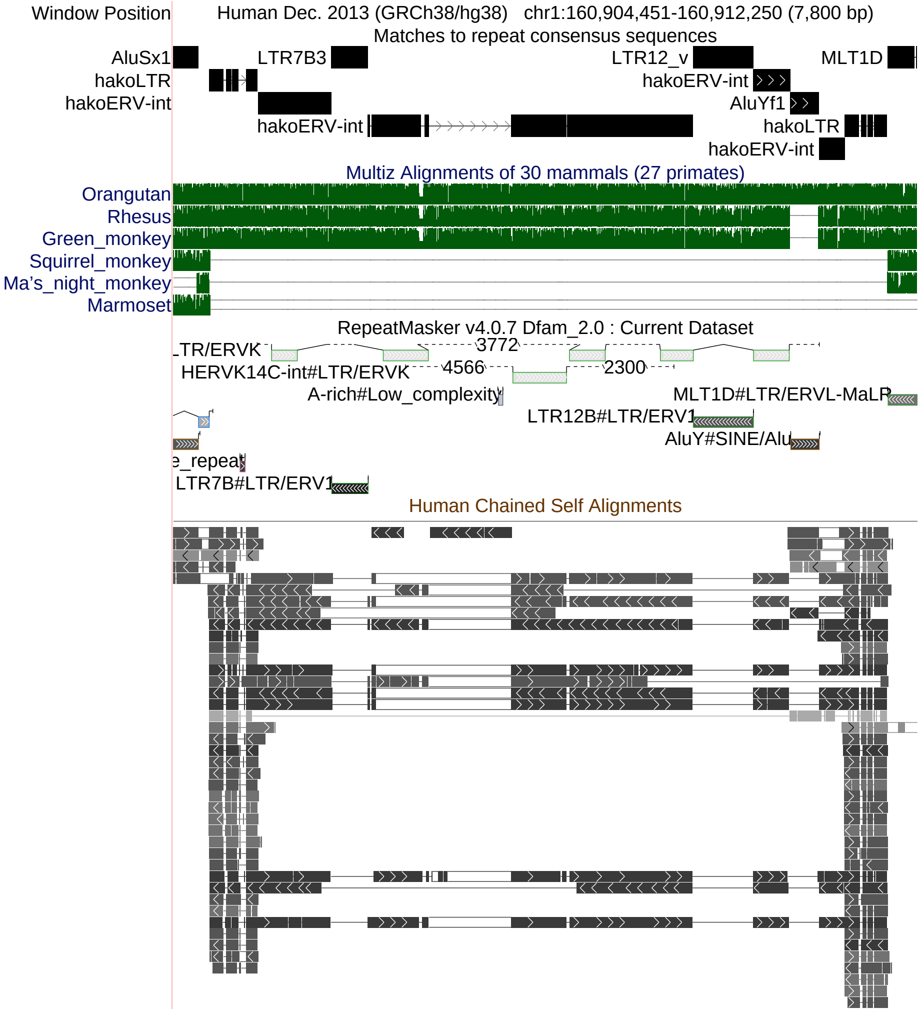


**Figure S3:** An ERV-Hako relic in human chromosome 1. Three other retrotransposons have inserted inside this ERV-Hako: an AluY element (gap in the rhesus and green monkey alignments), and two solo LTRs: LTR7B and LTR12. There is also a gap (arrowheads) in the match to the Hako consensus sequence, which coincides with one of the human chained self alignments. The DNA in this gap is homologous to part of the 3′-UTR of *SPHKAP*. Screenshot from http://genome.ucsc.edu.

```
Fig.1_left_LTR        tgttggggggtgcatggcaacaaattcaagcttgtatacaaagcatttgaggtcggggcat
hako_LTR_consensus    tgtggggggtgcacggcaacatattcaagcttatgtacaaggcatttgaggtcggggcat
hakoP_LTR_consensus   tgtagggggcgcacggcaacatattcaagcttatgtacatggcatttgaggtcggggcat

Fig.1_left_LTR        ggaaaaagacggaggcactgtgtgtatgttatttgtgcatggaaatgtaactccttgacc
hako_LTR_consensus    ggaaaaatactgaggcactgtgtgtatgttatttgtgcatgagaatgnaactccttgacc
hakoP_LTR_consensus   ggaaaaatactgaggcac--tgtgtatgttatttgtgcatgggaatgaaactccttgacc

Fig.1_left_LTR        ctgaaaacaggacaaggagtggagt-----gtgtcatgaggaacgctgaaaacagcctcc
hako_LTR_consensus    ctgaaaacaggacagggagtggagtgtgtggtgtgataaggaacgctgaaaacagcctcc
hakoP_LTR_consensus   ctgaaaacaggacagggagtggagtgtgtggtgtgataaggaacgctgaaaacagcctcc

Fig.1_left_LTR        tgagaatgtggtttgaatgcttttagaaggccacaggtgtgtcacgacccgacctcaaga
hako_LTR_consensus    tgagaatgcggtttgagtgcttttacaaggccacaggtgtctcacgacccgacctcaaaa
hakoP_LTR_consensus   tgagaatgcagtttgagtg--tgtacaaggtcacaggtgcctcagcaacccacctcaaat

Fig.1_left_LTR        ggccatctagtggatgtttgtagttta-acaagccctttcaataaatacttggcggatgg
hako_LTR_consensus    agccatctagtggatgtttgtggttta-acaagccctttcaataaatacttggcggacgg
hakoP_LTR_consensus   gaccatctagtggatgtttgtagttaatacaagccctttcaataaatacttggtggacgg

Fig.1_left_LTR        attctggggtgacactctctcagaagagtggtccgcc-gctccgctcagctggaattgtc
hako_LTR_consensus    atgctggggcgg-actctctcagaagagctgcccccc-gccccgctcagctggaattgtc
hakoP_LTR_consensus   atgctggggcgg-actctctcggaagagctaccccctagccccgctcagctggaactgtc

Fig.1_left_LTR        tgagaac-tcattctgggcgttcactgcaagctataaactctacа 397
hako_LTR_consensus    tgagaac-tcattcttggcgttcactgcaagctataagctctgca 401
hakoP_LTR_consensus   tgagtacttcgttcttggcgttcactgcaagctataagctctgca 400
```

**Figure S4:** An LTR in human chromosome 1 (the left LTR in Fig. 1) matches ERV-Hako rather than ERV-HakoP. The HakoP LTR consensus sequence differs from the other 2 sequences at 27 sites, and the Hako LTR consensus differs at 3 sites. Figure made with the Sequence Manipulation Suite [5].

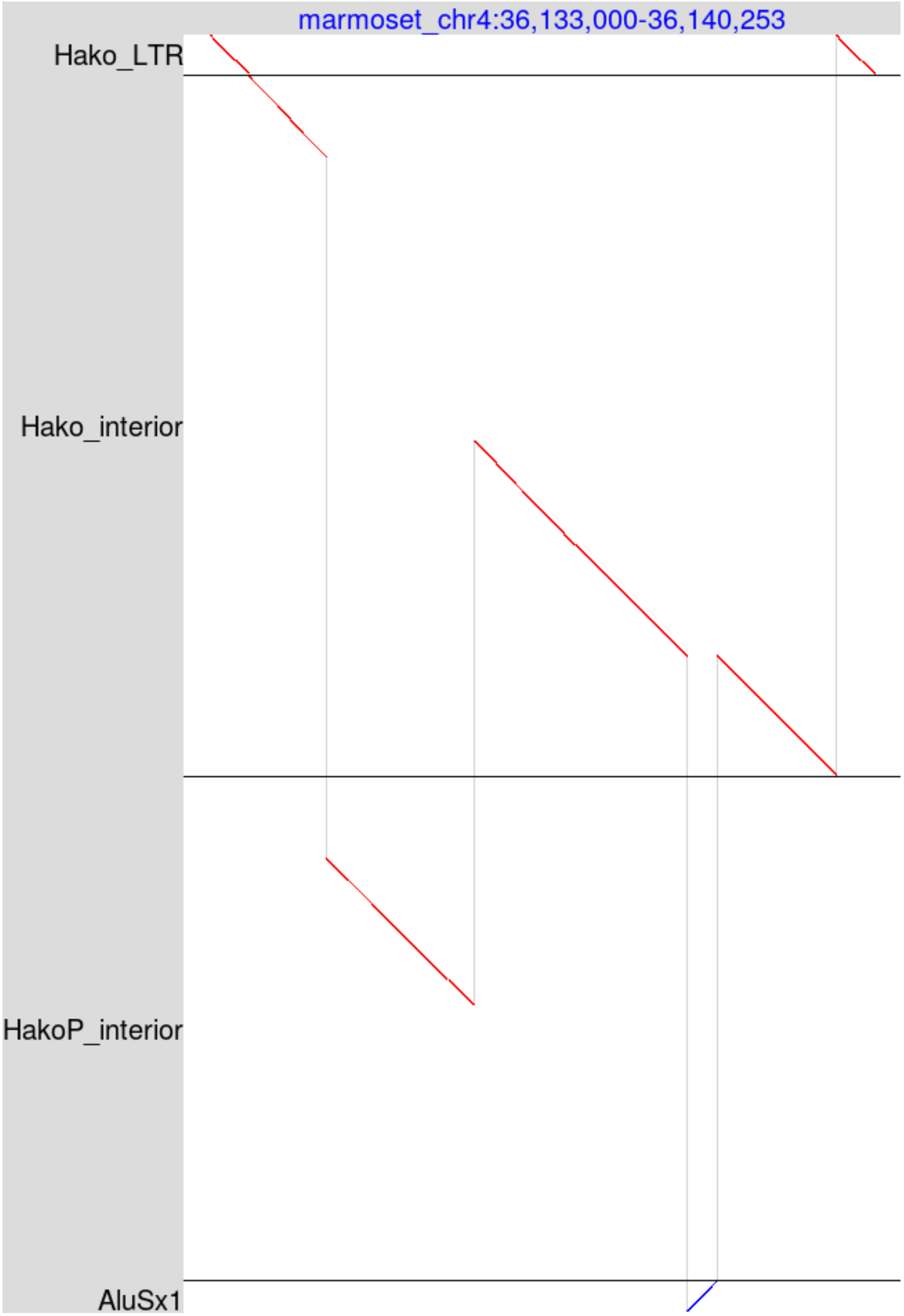


**Figure S5:** A defective ERV, related to Hako and HakoP, in chromosome 4 (NC_071445.1) of the marmoset *Callithrix jacchus*. This ERV is in the chromosome's reverse strand, so the chromosome is shown in reverse direction (indicated by blue color of the top chromosome label).

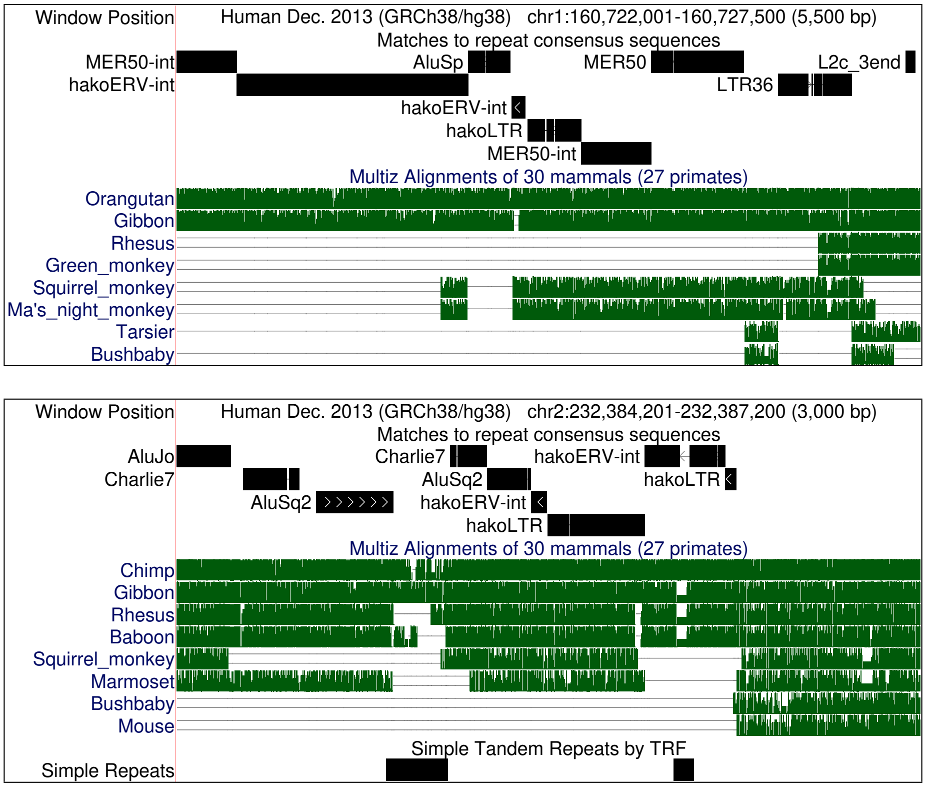


**Figure S6:** Two ERV-Hako relics, in human chromosome 1 (upper) and 2 (lower), that are common to catarrhines and platyrrhines.

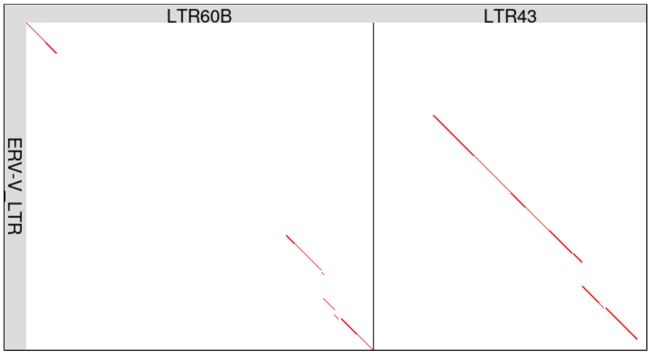


**Figure S7:** Parts of the ERV-V LTR are similar to two other kinds of LTR that are in the Dfam database.

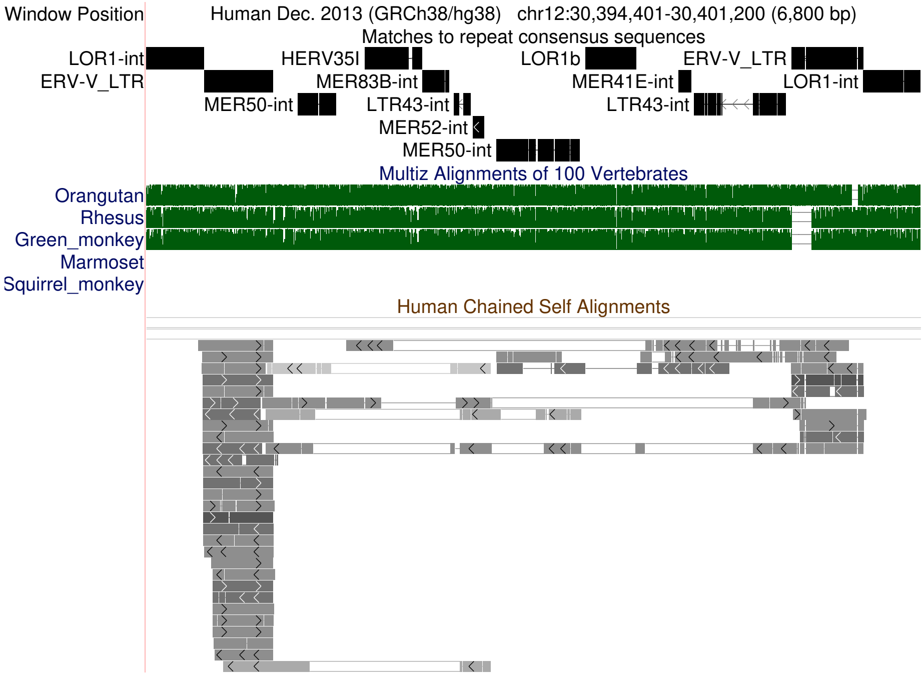


**Figure S8:** An ERV relic in human chromosome 12 that is flanked by ERV-V LTRs. The region between the LTRs matches various ERV consensus sequences in Dfam. LOR1b is a solo LTR inserted within this ERV.

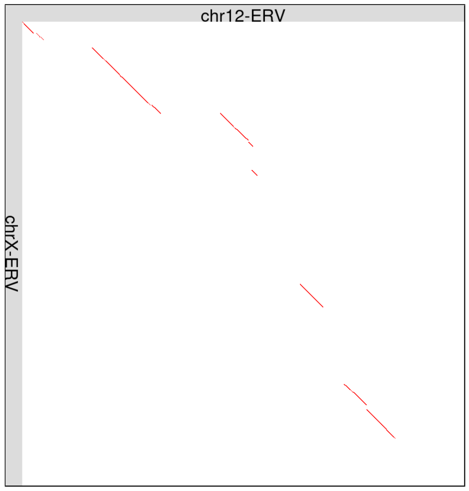


**Figure S9:** Comparison of two human ERV relics that have ERV-V LTRs. This figure just shows the internal ERV regions, excluding the LTRs. The ERVs are shown after removing retrotransposons (e.g. Alu) inserted within them.

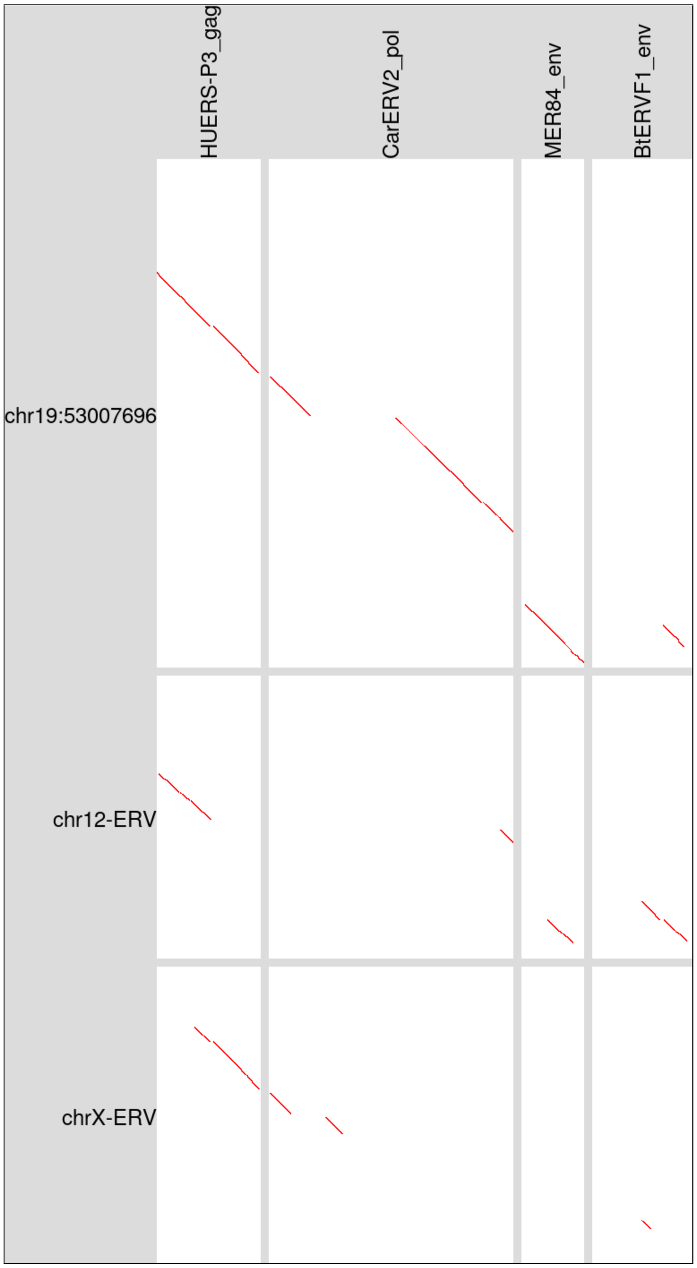


**Figure S10:** An ERV-V relic in human chromosome 19, and related ERV relics in chromosomes 12 and X, have DNA homologous to parts of gag, pol, and env genes. This figure was made by DNA-versus-protein alignment with LAST. The protein sequences (horizontal) are from RepeatMasker's file of protein sequences.

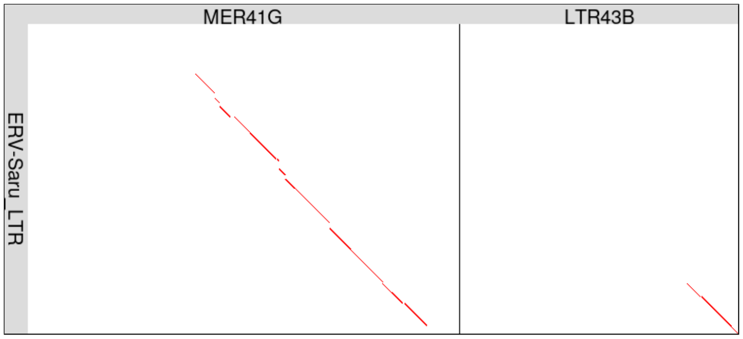


```
saruLTR 104 AGAAAGGAACTGTTACAACAGGCTGTAACAATGAAGACAGATTGCAGACATCCTGCTTTAGGCACAGATAAGAATGTAACAAAGAAGA---------AACTGGCCGC 201
            |||||||| |||||| ||   |||  ||  |  | |||           | ||||   |       |||||||   ||||||| | ||         | ||||| |
 MER41G 346 AGAAAGGAGCTGTTAAAATTAGCTACAAGGACAAGGACG----------AGCCTGGGCT-------GATAAGACCCTAACAAACAGGATGGGGGCTAAGCTGGCTGA 435

saruLTR 202 AACTGGTTAAATCCAAGATGGCCGAAAAACTTGACCGACTGCCGACCCTTGGCTTCATTATGCCCCTATTACCATA--AAATTTCCATGGGGAAGCCTCCCACTCCC 306
            ||| || |   ||||| |||||| |    | |||||| |   || ||||  | | | ||||| | |  ||||||||  ||||                | ||| |||
 MER41G 436 AACCGGCTGGGTCCAACATGGC-GCTGGATTTGACCCATGCCCTACCCCAGACCTAATTATACGCTCATTACCATACTAAAT----------------CACACACCC 525

saruLTR 307 ATCATGCACCTGACGCCATGACGGTTCCGGAT-TAACCATATTTAGTCAAGAAAAGGGTGGCACCCCGATTCCGGGAATTGCCCGCCCATTTCCCGGAAAACCCCTC 412
            | ||         ||||||||| | ||||    |  |||||||||||  | ||| |||||||||||| ||||   ||| | ||| ||  |||||  ||| |||
 MER41G 526 ACCA--------GCGCCATGACAGATCCGAGCATGCCCATATTTAGTATAAAAATGGGTGGCACCCCAATTCTAAGAAATCCCCNCCTTTTTCCTAGAANACC---- 620

saruLTR 413 CCCTTATTATAGAATATTCCGTGCCTTCATTATGCTTATCCATATAGTACGTGAGCCCTGACCACGTTGCGCGCGGCTCATTCTTTTGAGCGCGCCTGCACTCCTCT 519
                   |   || ||||||   || | ||||       ||||| | |  |  | |||  ||  ||||| ||||| |||  |||   |||| |||| ||||| ||||
 MER41G 621 -------TNATGATTATTCCACCCCCTAATTAGAAGAGCCCATAAAATTAG-AAACCCAAACTCCGTTGTGCGCGACTCGCTCTCNCGAGCACGCCCGCACTTCTCT 719

saruLTR 520 CTTGAGTGTGTGCTTGCTTTCGCTCCGCAAT-AAAGCTTCTATACTTTCACTGTG----GTCTCGCTTTCAAATTCTTTTGTGCGGCGAAGACAAGAACCTGAAC 619
            ||| ||||||    | ||||||||  ||||| |||||||||   ||||| ||       | ||||       ||||||||   ||| ||  | |||||||||| ||
 MER41G 720 CTTAAGTGTG----TACTTTCGCTTTGCAATAAAAGCTTCTTGCCTTTCGCTTCATTCTGACTCGTCCCTGAATTCTTTCTCGCGACGGTGTCAAGAACCTGGAC 820


saruLTR 534 TGCTTTCGCTCCGCAATAAAGCTTCT----ATACTTTCACTGTGGTCTCGCTTTCAAATTCTTTTGTGCGGCGAAGACAAGAACCTGAACTAGGCTTACTGGCAACA 636
            | ||||||||  |||||||| |||||     |||||| ||| ||| |||||| ||||||||||||||||||||||| |||||||||||||  |||  || |||||||
 LTR43B 468 TACTTTCGCTTTGCAATAAA-CTTCTTTGCCTACTTTTACTTTGGACTCGCTCTCAAATTCTTTTGTGCGGCGAAGTCAAGAACCTGAACC-GGCCCACCGGCAACA 572
```

**Figure S11:** Similarities between parts of the ERV-Saru LTR, and parts of two other LTRs in the Dfam database.

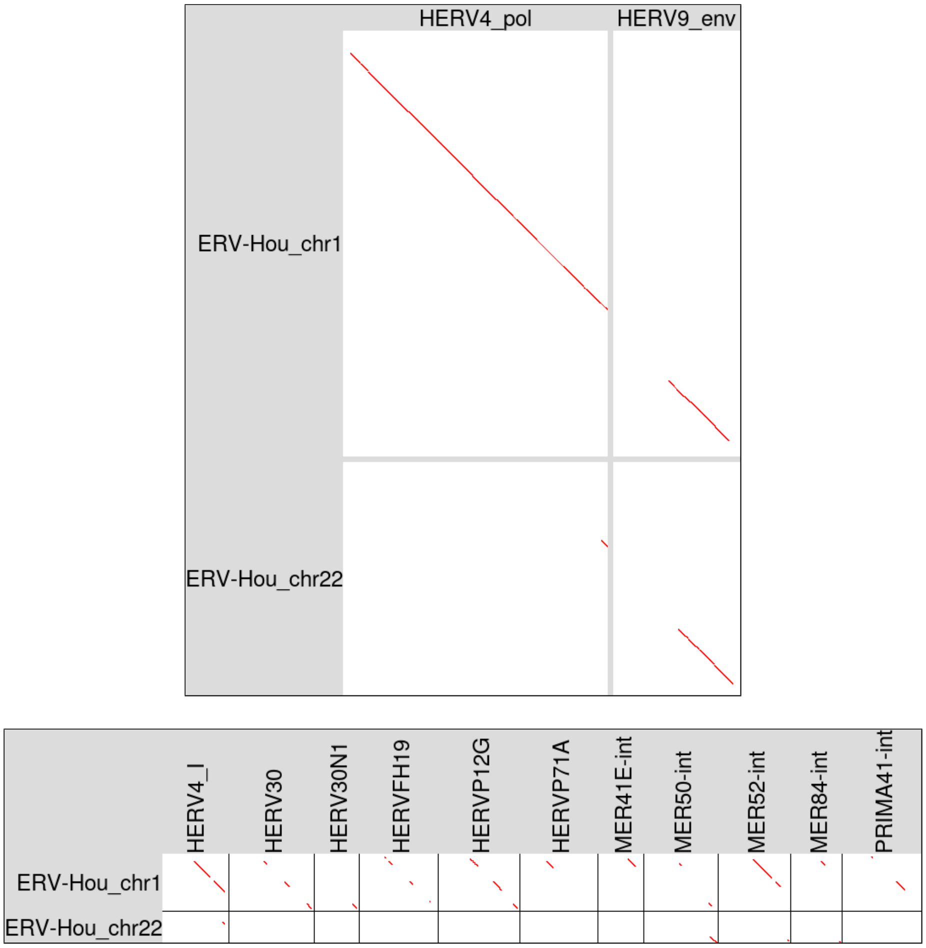


**Figure S12:** Homology of human ERV-Hou DNA sequences to: ERV proteins (above), and ERV DNA consensus sequences in Dfam (below). All these DNA sequences are internal ERV regions excluding LTRs. The chr1 ERV-Hou is shown after removing a LINE fragment that was recently inserted inside it. The DNA-to-protein alignments were made with LAST: the proteins (HERV4_pol and HERV9_env) are from RepeatMasker's file of protein sequences.

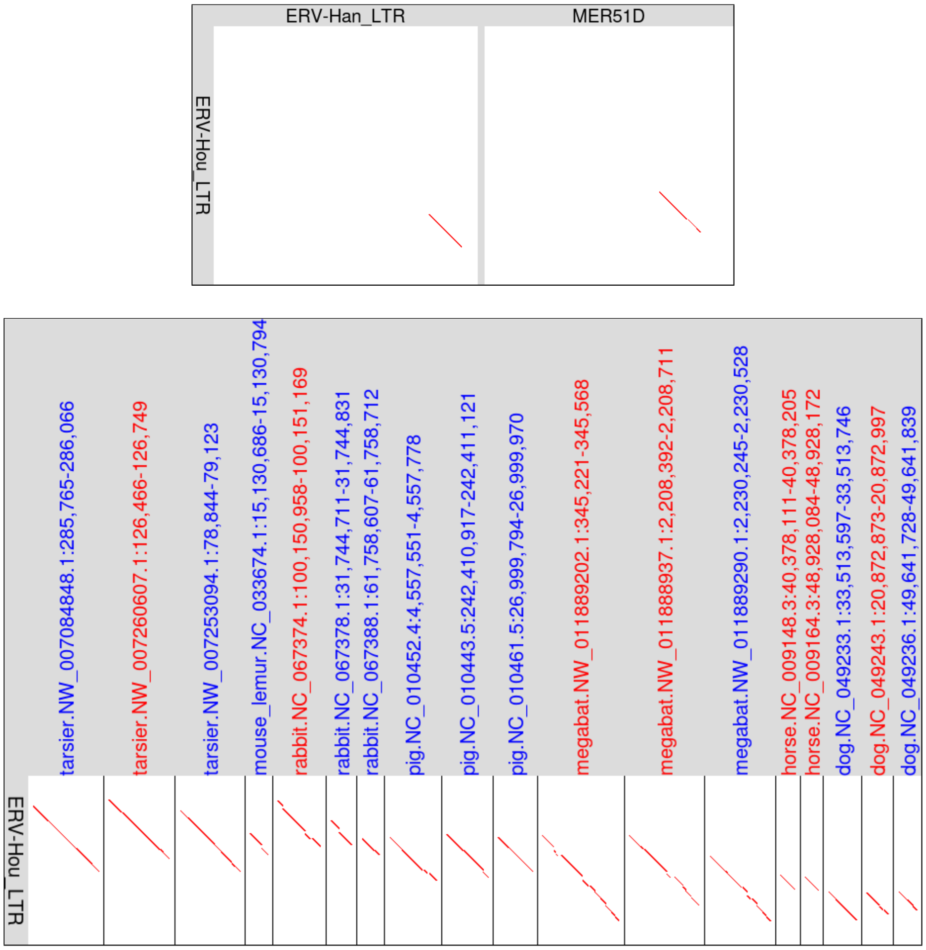


**Figure S13:** Upper: partial homology between ERV-Hou LTR and other types of LTR. Lower: homology to parts of ERV-Hou LTR in various mammal genomes. Blue labels indicate that the reverse DNA strand is shown; red labels indicate forward strand.

```
human.chr12 18467644 TGTTAGGTAGTGAGATAGACATTAGCAGCTGGGAAGGGGTAAGAGAAGAGAGCAGAAAAGCCATCTCTAAAACTGCATCTGGCCCACCTAAGTTCAGCTCTGGAACCACCCTAACTC 18467760
                     ||||||||||| |||||||||||||||||||||| ||||| |||||||||||||||| ||  || |||| || ||  |||||||||||||||||||| |    ||| |||||||||
ERV-Hou_LTR 1        TGTTAGGTAGTTAGATAGACATTAGCAGCTGGGAGGGGGTGAGAGAAGAGAGCAGAAAGGCTGTCACTAAGACAGCCCCTGGCCCACCTAAGTTCAGCCCCAAGACCGCCCTAACTC 117

human.chr12 18467761 CACCCTAACAGATGGAGTTTTTGGTGGAGACTGTATCCAGCACATCTTATAGAAAGAGAAACTAGAGCACAAGTGGAAATCCCCCAAAGTAGCACATGCCCAATAACCTAAAGCTCT 18467877
                     ||||||||| |||||||||| ||||  || ||||  ||||||||||| |  |||| |||||||||| |||||| ||||||||||||| ||||| ||||||||||| |||||||||| || |
ERV-Hou_LTR 118      CACCCTAACGGATGGAGTTTGTGGTAAAGTCTGTGGCCAGCACATCCTGGAGAAGGAGAAACTAGGGCACAGGTGGAAATCCCCTAAAGTGGCACATGCCCAGTAACCTAAAACTGT 234

human.chr12 18467878 ATCTTGGAATTGACCCTAGGCTCATGATACCATTATTATAATAAAATTTACATGTGGTTTTTGCT--CCCCTGAGTGGGCATTGTTTTAAAACAAATTATAGGTAAAAATATGCACA 18467992
                     ||| |    || |||| | |||||| ||||||| ||||||||||||||||||||||||||||  |   |||| |||||||| ||  ||   ||| |||||| |||||||| |||| ||
ERV-Hou_LTR 235      ATCCTCAGGTTNACCCCAAGCTCATTATACCATCATTATAATAAAATTTACATGTGGTTTTNTCCTGCCCCCGAGTGGGCTTTTCTT---AACGAATTATGGGTAAAAACATGCGCA 348

human.chr12 18467993 GTTTAATTTTAGCTACATAAACATAAACTGCCAATCAAATGACATCATCCTGTCACTCAAACACAGCCCAAACCTCAACTGCTCCCCAGAAA-CCCATAAAAGGACCTCAAGTTTTG 18468108
                     |||||||||||| || |||| ||||||||||||||||||||||||||||||||||||||||| |||||||||||||||||||| ||||||| ||| |||||||||| |||   || | ||
ERV-Hou_LTR 349      GTTTAATTTTAGTTATATAACCATAAACTGCCAATCAAATGACATCATCCTGTCACTCAGACACAGCCCAAACCTCAACTCCTCCCCACAAACCCCATAAAAGCACCCTGAGCTCTG 465

human.chr12 18468109 TAAAGAGGTGCTGATCTCACTTCACAAAAATAAGTCTGCTCTCCCTCTGAGAGTATATTTCTATGCTTCAATTAACTTTGCTTTAAGCTTGCATTCTGGAGTTAGTCTGCAATTTTT 18468225
                     |||||||| |||||| ||||||| || |||| || | |||||||||||||||| |||| || ||||||||| |||||||||| ||||||||| ||| |||||| ||||| || |
ERV-Hou_LTR 466      TAAAGAGGGGCTGATTTCACTTCGCAGAAATCAGCCCGCTCTCCCTCTGAGAGTGTATTACTGTGCTTCAATAAACTTTGCTTTGAGCTTGCATTTTGGTGTTAGTTTGCAA-TTCT 581

human.chr12 18468226 TTGTTTACTCTCACAAGAACTGAGATTGCTGGTCCAGAGCTCCAACTCTGTTCATCTCCTTGGTTAAAGAATCCATTCCAATGCAGAATTCCCACTGACA 18468325
                     ||| | ||| |||||||||| ||||||||||||||||||||||||  ||||||| ||||||| |||||||| |||||| |||| |||||||||||  | |||
ERV-Hou_LTR 582      TTGCTCACTATCACAAGAACCGAGATTGCTGGTCCAGAGCTCCGGCTCTGTTGATCTCCTCGGTTAAAGGATCCATCCCAACGCAGAATTCCCGGTAACA 681


dolphin.chr15 48140533 TGTTAGGCAGTTAGTCAGACATTAGCAGCAAGGAGGCCACAGTGGCAAAAGACAAGAGGGGAGTAGTCAAGTTACACCC--ACTCGTCTGGG---------GACCCTCCCCTGACCC 48140638
                       ||||||| ||||||  ||||||||||||  |||||        | | |   ||  | ||  ||    |||  |  |||   | |  ||   |          |  |  |||| || |
ERV-Hou_LTR   1        TGTTAGGTAGTTAGATAGACATTAGCAGCTGGGAGGGGGTGAGAGAAGAGAGCAGAAAGGCTGTCACTAAGACAGCCCCTGGCCCACCTAAGTTCAGCCCCAAGACCGCCCTAACTC 117

dolphin.chr15 48140639 CACCCAGAATGGGCAAAACTATGGTCATGT-GGTGGGCAATTTATCCTGATAAAGAGGGAACACAGTAACACTAGGGAACTTCCCTGGGCTGCGCATGCCCAGACTAGACAGGTGTA 48140754
                       |||||  |  |    |   | |||| | ||  |||| ||    ||||||   |||  | |||   |  |||   || ||   ||    |  || |||||||||
ERV-Hou_LTR   118      CACCCTAACGGATGGAGTTTGTGGTAAAGTCTGTGGCCAGCACATCCTGGAGAAGGAGAAACTAGGGCACAGGTGGAAATCCCCTAAAGTGGCACATGCCCAG-------------- 220

dolphin.chr15 48140755 TAACCTATAG--------------TAAAACCAAACTCATTATACCATCACTATAATAAAATATGCATATGGTTTGGCAAACCGCCCCCCACCCCGCTGACCCCG----GGCTGTTCT 48140853
                       ||||||| |                   | |  |||| ||||||||||||||| ||||||||||| | ||| ||||||                        ||| |||||    |||| ||||
ERV-Hou_LTR   221      TAACCTAAAACTGTATCCTCAGGTTNACCCCAAGCTCATTATACCATCATTATAATAAAATTTACATGTGGTTT-----------------TNTCCTGCCCCCGAGTGGGCTTTTCT 320

dolphin.chr15 48140854 TAGCGAATCAAGGGTAAGGGCATGAGCAATAATGAACTTGTTACATAGATGGGGACTGTCAATCAAATAATGATACCCCTGTCACTCACCCATGATT-GCGCCCCGCCTTCTCCCAG 48140969
                       || ||||| | ||||||   |||| ||| |       | |||| |||       |||| ||||||||| |    | || |||||||||  ||         || |  || |||||
ERV-Hou_LTR   321      TAACGAATTATGGGTAAAAACATGCGCAGTTTAATTTTAGTTATATAACCATAAACTGCCAATCAAATGACATCATCC-TGTCACTCAGACACAGCCCAAACCTCAACTCCTCCCCA 436

dolphin.chr15 48140970 AACTCTCCTTATAAACACCTTTAAGCCTGTACAGGAGAGCTGGTTTCACTTAGTAGAAA-CAGCCCACTCTCCCTCTGAGAGTGTA--ACTATACTTTAATAAAACTTTGCTTTGCA 48141083
                        |  | || || || |||| | |   ||||| ||  | |||| |||||||| | ||||| |||||| ||||||||||||||||||  ||| | ||| ||| ||||||||||||
ERV-Hou_LTR   437      CAAACCCCATAAAAGCACCCTGAGCTCTGTAAAGAGGGGCTGATTTCACTTCGCAGAAATCAGCCCGCTCTCCCTCTGAGAGTGTATTACTGTGCTTCAAT-AAACTTTGCTTTGAG 552

dolphin.chr15 48141084 CTTAAGCCCAAAGTGGTAGTCTGCAATTCTTTGCTTGCTCTCACAAGGACCGAGATTGCTGGTCCGG-----------CATTGACCTCCTCTGTT---GGAGCTATTCTGACACA 48141184
                       |||         ||| |||| |||||||||||||||  || ||||||| ||||||||||||||||| |                |||| |||||| |||   ||| | || |  || ||
ERV-Hou_LTR   553      CTTGCATTT-TGGTGTTAGTTTGCAATTCTTTGCTCACTATCACAAGAACCGAGATTGCTGGTCCAGAGCTCCGGCTCTGTTGATCTCCTCGGTTAAAGGATCCATCCCAACGCA 666
```

**Figure S14:** Above: a Hou LTR relic in the human genome (84% identity). Below: a Hou-like LTR relic in the dolphin genome (59% identity).

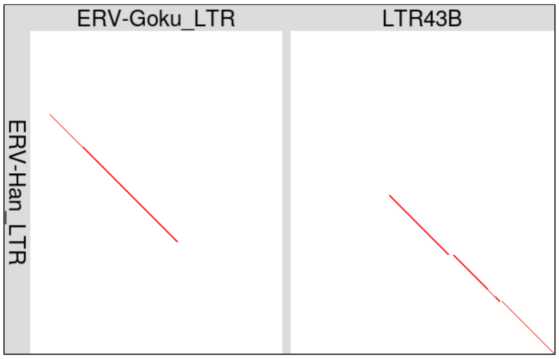


**Figure S15:** Parts of the ERV-Han LTR are similar to other LTRs.

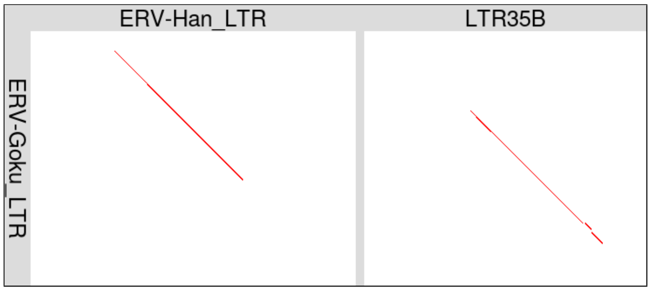


**Figure S16:** Parts of the ERV-Goku LTR are similar to other LTRs.

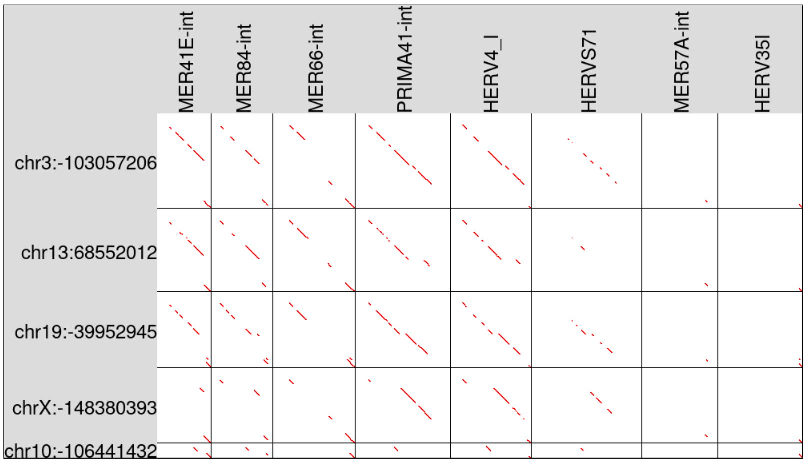


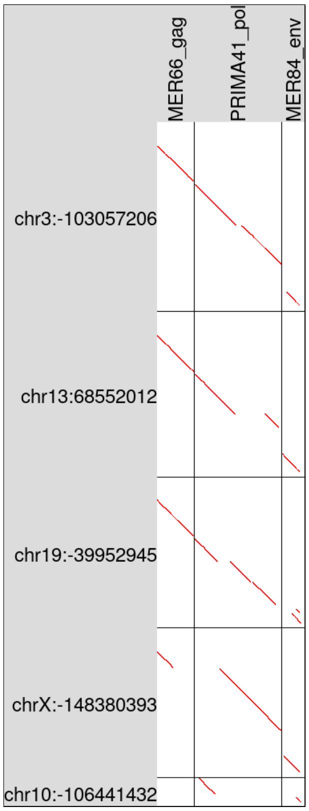


**Figure S17:** Homology between five ERV-Goku internal regions in human genome hg38 (vertical), and: ERV consensus DNA sequences (upper), and ERV proteins (lower). The five Goku sequences are shown after removing younger insertions within them. They are labeled by chromosome and 5′ coordinate: "-" indicates reverse strand.

```
human.chr12  4807220 tgagatgggcaagcTGGCTGCTTTCCTGTTGTGATGTGGTTTGGAGAAAAAAGAACAAAAGCCCTTTACTCAAGCTGTA---------ACTCCCAGCCAGTCAGCATCAAAG 4807322
                     |||||  |||| |||||||| || |||||| ||||||||| ||||| |||| || ||||||||| ||||||||||| ||         |||| |||||| |||||| ||||
ERV-Goku_LTR 1       TGAGACaggcaggctggctggtTCCCTGTTTTGATGTGGTCTGGAGGAAAAGGA-CAAAAGCCCCTTACTCAAGCTNTAGCTTATCTAACTCTCAGCCAATCAGCAACAAAA 111

human.chr12  4807323 G-CCCAAGAAGCTATTAACCACAAATTAATTAACCCCTTGCTTTAGAGAACTAAGGACCTTTCTGAGGCCCTACGTGCCTAGCTAGGCTTAACTTTCACCTCATCATGAACT 4807433
                     | |||||||||||||||||| ||| ||         | ||||| || |  ||| |||| |  | |  ||||  | |||  || ||| |||||  | || |  ||  | | |
ERV-Goku_LTR 112     GACCCAAGAAGCTATTAACCGCAAGTT---------CCTGCTTCAGGGGGCTAGGGACTTCCCCGGAGCCCCGCATGCGCAGTTAGACTTAAACTCCAACCTATAGTTACCC 214

human.chr12  4807434 TTTCCTTATTTTAGTACTAAAAATCATGCCCACAGGTGGAGATTTAAGATGCTAATGAGACATATGTTGTATGAAGCAGCATCTTAAGCCACCGTACATGTGCCCGAAAAAC 4807545
                      ||||| |||||| | |||||||||| |||||  ||||||||||||| |||||||||  ||||  | ||||||||| ||||| || |||||| |  || | ||  |||||||
ERV-Goku_LTR 215     CTTCCTCATTTTAATGCTAAAAATCACGCCCAGGGGTGGAGATTTAAAATGCTAATGCTACATGCGATGTATGAAGAAGCATGTTGAGCCACTGCGCAAGCGCTAGAAAAAC 326

human.chr12  4807546 CTCACCTCTACATGCCCTG-----ACTTCCCCTTACTGCAGACCTCCACAAAGGGAACCCACACCTTGACTTTGGAGAGCAACCCA-------CTTCCTTTCTTGGTGTTTG 4807645
                     | | ||| |||||||||||     ||    || ||  | |   | | | |||   |||||||||  |  | | || ||||| ||||        ||||||||| |||| | |
ERV-Goku_LTR 327     CCCTCCTATACATGCCCTGACGTAACCCTTCCCTATAGAAAGACCCTATAAAACTAACCCACACACTACCCTCGGGGAGCAGCCCATTCCCTTTTTCCTTTCTCGGTGCTGG 438

human.chr12  4807646 GTCCCTTATGTCCATAAGCTTTCATAAAATCTTTCTCTTTGCTACTGTATGCTGTGATCTCTTTTGATTTCTATCCTGGGGGATCAAAAAAGCCCACAGGGCGTTGGTGCCA 4807757
                      |||||| ||  || |||||   | | || ||||| ||||||| || | |   ||||||||| ||||||||||||||||| ||||  || | ||  |||||||  |||  ||
ERV-Goku_LTR 439     CTCCCTTGTG--CACAAGCTNAAAAATAAACTTTC-CTTTGCTGCTATGTCTGGTGATCTCTCTTGATTTCTATCCTGGGAGATCGCAAGAACC--CAGGGCGCCGGTAACA 545


lemur.chr10  65589990 TGAGACAGGCAGACTGGTTGGTTTCCTATTTTGGTGTGGTCCACAC-AAGGGAGCAGAGGTCTCTTGCTCAAGCTGAGGACCTCTGAATTTCTAGCCAATTAGAATCAAGAG 65590100
                      |||||||||||| |||| ||||| ||| ||||| |||||||   |  ||  |  || | | | ||| ||||||||   |       || |   ||||||| || | ||| ||
ERV-Goku_LTR 1        TGAGACaggcaggctggctggtTCCCTGTTTTGATGTGGTCTGGAGGAAAAGGACAAAAGCCCCTTACTCAAGCTNTAGCTTATCTAACTCTCAGCCAATCAGCAACAAAAG 112

lemur.chr10  65590101 ACC-AAAAACCTGTTAACCACAAGTTTCTGTTGTAGAGGGCTAGGAACTTTCCTGGAGCCTTTACAGGTGCAGTTAGGCTTAACCTTCAAATTACAGTAACACTTTCTTCAT 65590211
                      ||| || || || |||||| |||||| ||| |  || |||||||| |||| || ||||||    || | |||||||| ||||| || |||  || ||| || | ||| ||||
ERV-Goku_LTR 113      ACCCAAGAAGCTATTAACCGCAAGTTCCTGCTTCAGGGGGCTAGGGACTTCCCCGGAGCCCC-GCATGCGCAGTTAGACTTAAACTCCAACCTATAGTTACCCCTTCCTCAT 223

lemur.chr10  65590212 TTTAATAGTAAAAATCACACCCAGAAGTGGAGATTTAAAATGCTAATGTTA-ATACAATACAT---GGAGTATGTAAAGCTACTGTGCAGGCACAAGAAAAACCCCTCCTAT 65590319
                      ||||||  |||||||||| |||||  |||||||||||||||||||||| || || | ||  ||   | || ||||  ||| |||| ||| || | |||||||||||||||||
ERV-Goku_LTR 224      TTTAATGCTAAAAATCACGCCCAGGGGTGGAGATTTAAAATGCTAATGCTACATGCGATGTATGAAGAAGCATGTTGAGCCACTGCGCAAGCGCTAGAAAAACCCCTCCTAT 335

lemur.chr10  65590320 ACATACTTTGACATAACC----------------------------AATTCCCACACAATTCTTGAGGAGCAGCCCAAACTCTTCTT-----------------TTCCTTAT 65590386
                      |||| |  |||| |||||                            ||  | ||||| |  || | |||||||||||  | ||| ||                 | |||| |
ERV-Goku_LTR 336      ACATGCCCTGACGTAACCCTTCCCTATAGAAAGACCCTATAAAACTAACCCACACACTACCCTCGGGGAGCAGCCCATTCCCTTTTTCCTTTCTCGGTGCTGGCTCCCTTGT 447

lemur.chr10  65590387 GTACCAGGTAAATAAAACGTTTTTCTTCATTGCTATGTCTGGTGATCTCTCTTGATTTCTATCCTGGGAGAGTTACAAGAACACAGGAGCATCAGTAACA 65590486
                      | || || | || || |   ||| |||   ||||||||||||||||||||||||||||||||||||||||| |  ||||||| |||| ||  | ||||||
ERV-Goku_LTR 448      GCACAAGCTNAAAAATAAACTTTCCTTTGCTGCTATGTCTGGTGATCTCTCTTGATTTCTATCCTGGGAGA-TCGCAAGAACCCAGG-GCGCCGGTAACA 545
```

**Figure S18:** Above: an ERV-Goku LTR relic in the human genome (70% identity). Below: an ERV-Goku LTR relic in the mouse lemur genome (68% identity).

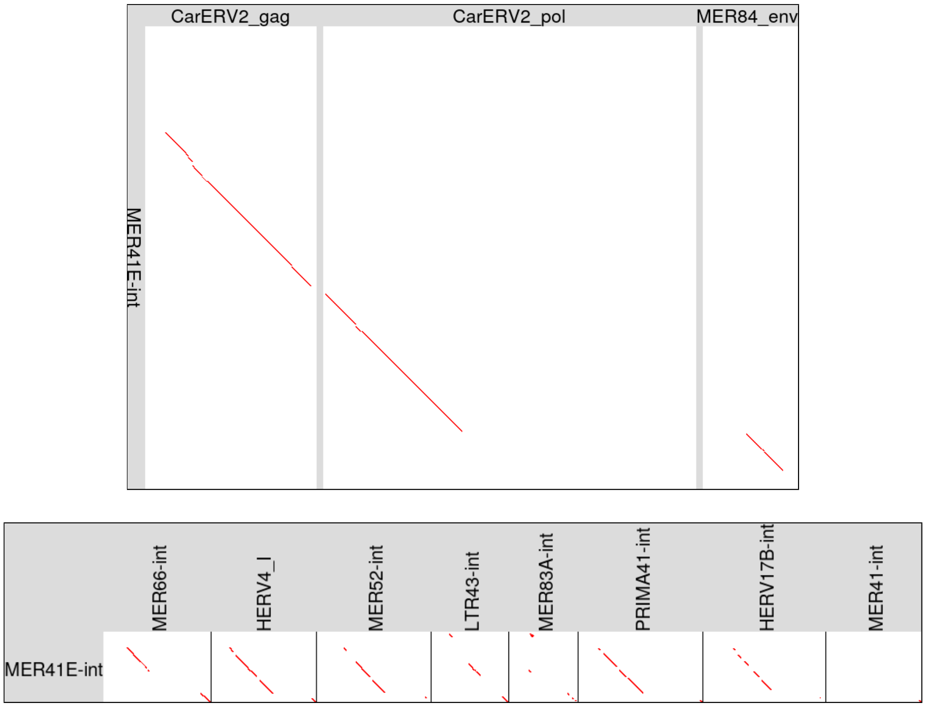


**Figure S19:** Homology of the MER41E DNA consensus sequence to: ERV proteins (above), and ERV DNA in Dfam (below). All these DNA sequences are internal ERV regions excluding LTRs.

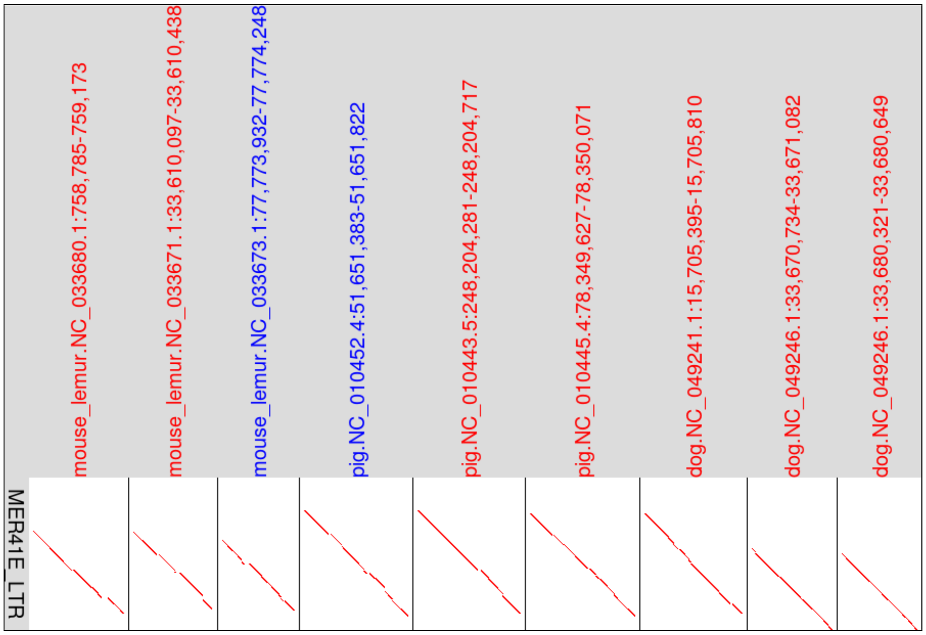


**Figure S20:** Homology to parts of ERV-MER41E LTR in various mammal genomes. Blue labels indicate that the reverse DNA strand is shown; red labels indicate forward strand.

```
human.chr15  51652211 TGAGAGAGGAATAATATAAGGTGGCCGTAGGAGATTAGAAAATTCCAGGCAACAATTTCATATGACTAGCAAAAGGCAACTGTTGAAATAGCTGCAGAGGCTATGGGCTGATAAGACCCT 51652330
                      ||||| ||||||||||| | |||| || |||| |||||||||||||||| || ||||| ||||||||||||||| ||||||||||||||||||||| |||| ||||||||||||||||
MER41E_LTR   1        TGAGACAGGAATAATACAGGGTGGTCGCAGGAGAATAGAAAATTCCAGGCAGCAGTTTCACATGACTAGCAAAAGGAAACTGTTGAAATAGCTGCAGAAGCTAGGGGCTGATAAGACCCT 120

human.chr15  51652331 GAGAAAC-AGGGTGTGGACCAAGATGGCTAAGACTGACTGAACCCAACATGTCACTGGATTTGACGTAGGTTTCTCCTAGAACCTCATTATATGCTCATTAACGTA----ATCACATGCC 51652445
                      || |||| ||||||||| ||||| |||||||||| ||||| |||||||||| | |||||||||||| ||||||||| ||||| ||||||||||| |||||||||| ||    ||||||  ||
MER41E_LTR   121      GAAAAACCAGGGTGTGGGCCAAGCTGGCTAAGACCGACTGGACCCAACATGGCGCTGGATTTGACCTAggtttcacctaggACCTCATTATACGCTCATTAACATACTAAATCACACACC 240

human.chr15  51652446 CACCAGCACCATGACACTTCCAGGAACACCCATTATTTAGTGCAAAAGTGGGTGGCACCACAGTTTTGAGAAATCTTCACCTTTTTCCAGGAATCTTCATGAATATGTCACCCCTTGGTT 51652565
                      ||||||| |||||||| |||| ||||||||||| |||| ||| |||| |||||||||||||||||  ||||||||| ||||||||||||||||||||||||||||||||  ||||||||||||
MER41E_LTR   241      CACCAGCGCCATGACAGTTCCGGGAACACCCAT-ATTTGGTGTAAAAATGGGTGGCACCACAGTTCCGAGAAATCTCCACCTTTTTCCAGGAATCTTCATGAATATTCCACCCCTTGGTT 359

human.chr15  51652566 ACAGAAACCCATAAAGATAGTACCCGAAAACC---TTGTGTGCAACTC---TCTTGAGTACACCTGCACTCCCCTTTCTTGAGTGTGTACTTTTCGCTTTGCAATAAATCTTCATCTTTT 51652679
                      | |||||||||||||| ||| | ||  |||||   ||| | || ||||   |||||||||| || |||||||||||||||||||||||||||||||||||||||||||||| | |  |||
MER41E_LTR   360      AAAGAAACCCATAAAGGTAGAAGCCCCAAACCCCNTTGNGCGCGACTCCTCTCTTGAGTACGCCCGCACTCCCCTTTCTTGAGTGTGTACTTTTCGCTTTGCAATAAATCTCCGTACTTT 479

human.chr15  51652680 CACTATTTTCTGACTCATCCCTGAATTTCTTCTTGTGATGGTGTCAAGAGCCTGGATACCGGTTGAGGTCAAGGTCCCACTAGTGTTTGTGGACCTACCCCAGCCCATCAGTATCA 51652795
                      |||||||||||||||| ||| |||||| ||||| | || |||||||||||||||||| ||||| || |||| |||||||||  | ||||| |||||| |||||||||| | ||||||
MER41E_LTR   480      CACTATTTTCTGACTCGTCCTTGAATTCCTTCTCGCGACGGTGTCAAGAGCCTGGACACCGGCTGGGGTCGAGGTCCCACCGGCGTTTGGGGACCTCCCCCAGCCCACCGGTATCA 595


tarsier.Scaffold91532 675157 TGAGATAGGAATAGTATTGGGTGGCCATGGAGGATGGGAAAATAT--GGCAGCAGTTAAGAAATCACATGACCAGTCAAGGAAGTACTATTACAACCAGCTAC 675257
                             ||||| ||||||| ||  |||||| |   |  ||   |||    ||||||||||      ||||||||| ||  || | |  ||| ||  ||  |||| |
MER41E_LTR            1      TGAGACAGGAATAATACAGGGTGGTCGCAGGAGAATAGAAAATTCCAGGCAGCAGTT------TCACATGACTAGCAAAAGGAA-ACTGTTGAAAT-AGCTGC 95

tarsier.Scaffold91532 675258 AAAGAC--AAAGCCCATAATGCCCTGAAAAAACAGGAGGTAG-TCAAACCAGCTAAGGCTAGCTGGATCCAACATGGAAATGGACGTTGACTTGCAAGCGACC 675357
                             | |  |     ||  ||||  |||||||||| ||||  || |  ||| |  |||||| |   ||||| |||||||||   ||||  ||||| |       |||
MER41E_LTR            96     AGAAGCTAGGGGCTGATAAGACCCTGAAAAACCAGGGTGTGGGCCAAGCTGGCTAAGACCGACTGGACCCAACATGGCGCTGGAT-TTGACCTAggtttcacc 197

tarsier.Scaffold91532 675358 TCAGGGCTCATTATACCCTCCTTAGCGTG-------ACACACCCCCTAGCACCATGACGGTTCCAGTTTTGATTAAATAAGGACAGAAAAGGGGAGGGTACTC 675453
                             |  |  |||||||||| ||| ||| | |        |||||||| | ||| ||||||| ||||| |        | ||  ||    |||| ||| ||   | |
MER41E_LTR            198    taggACCTCATTATACGCTCATTAACATACTAAATCACACACCCACCAGCGCCATGACAGTTCCGGGAACACCCATATTTGGTGTAAAAATGGGTGGCACCAC 300

tarsier.Scaffold91532 675454 AATTCCAGGAAATCTTCACATCTGCCTCAGAAAACCCGTGAATATTCCACCCCATAGTTAGAGATAGCTATAAAGAGAAACACCCCAAACCCCACTGGGGCTT 675556
                             | ||||  ||||||| ||| | |  |   |||    | ||||||||||||||| | |||| |||   | ||||||  | |  ||||||||||||  || |
MER41E_LTR            301    AGTTCCGAGAAATCTCCACCTTTTTCCAGGAATCTTCATGAATATTCCACCCCTTGGTTAAAGAAACCCATAAAGGTAGAAGCCCCAAACCCCNTTGNG---- 399

tarsier.Scaffold91532 675557 GAGTATACTCATATTCTCTTGAGTATGCTCATACTCCCTCTCTCTTGAGTATATACTTTT-GCTTTACAATAAGCCTCCTTTGCTTTGACTTTACTCTGACTC 675658
                                    |   |  ||||||||||| || |  |||||| || |||||||| | ||||||| ||||| ||||||  ||||  | |||| ||| |  ||||||||
MER41E_LTR            400    -----CGCGACTCCTCTCTTGAGTACGCCCGCACTCCC-CTTTCTTGAGTGTGTACTTTTCGCTTTGCAATAAATCTCC-GTACTTTCACTATTTTCTGACTC 495

tarsier.Scaffold91532 675659 ATCCTTGAATTCTTTCTCATAAAGATGTCAAGAACTTGGCTTCCCAGGTGAGGTCGAGCGTCTCCCCTGCTTCAGGAGATCTTTCCAAGCCCTCTGACATTA 675760
                              ||||||||||| |||||   | | |||||||| | |||    || | || ||||||| ||| | || || |  || || ||  ||||| | |  || |
MER41E_LTR            496    GTCCTTGAATTCCTTCTCGCGACGGTGTCAAGAGCCTGG-ACACCGGCTGGGGTCGAG-GTCCCACCGGCGTTTGGGGACCTCCCCCAGCCCACCGGTATCA 595
```

**Figure S21:** Above: a MER41E LTR relic in the human genome (86% identity). Below: a MER41E-like LTR relic in the tarsier genome (65% identity).
